# Regulation of Hepatic Xenosensor Function by HNF4alpha

**DOI:** 10.1101/2023.10.11.561888

**Authors:** Manasi Kotulkar, Diego Paine Cabrera, Dakota Robarts, Udayan Apte

## Abstract

Nuclear receptors including Aryl hydrocarbon Receptor (AhR), Constitutive Androstane Receptor (CAR), Pregnane X Receptor (PXR), and Peroxisome Proliferator-Activated Receptor-alpha (PPARα) function as xenobiotic sensors. Hepatocyte nuclear factor 4alpha (HNF4α) is a highly conserved orphan nuclear receptor essential for liver function. We tested the hypothesis that HNF4α is essential for function of these four major xenosensors.

Wild-type (WT) and hepatocyte-specific HNF4α knockout (HNF4α-KO) mice were treated with the mouse-specific activators of AhR (TCDD, 30 µg/kg), CAR (TCPOBOP, 2.5 µg/g), PXR, (PCN, 100 µg/g), and PPARα (WY-14643, 1 mg/kg). Blood and liver tissue samples were collected to study nuclear receptor activation.

TCDD (AhR agonist) treatment did not affect the liver-to-body weight ratio (LW/BW) in either WT or HNF4α-KO mice. Further, TCDD activated AhR in both WT and HNF4-KO mice, confirmed by increase in expression of its target genes. TCPOBOP (CAR agonist) significantly increased the LW/BW ratio and CAR target gene expression in WT mice, but not in HNF4α-KO mice. PCN (a mouse PXR agonist) significantly increased LW/BW ratio in both WT and HNF4α-KO mice however, it failed to induce PXR target genes in HNF4 KO mice. The treatment of WY-14643 (PPARα agonist) increased LW/BW ratio and PPARα target gene expression in WT mice but not in HNF4α-KO mice.

Together, these data indicate that the function of CAR, PXR, and PPARα but not of AhR was disrupted in HNF4α-KO mice. These results demonstrate that HNF4α function is critical for the activation of hepatic xenosensors, which are critical for toxicological responses.

## Introduction

The xenobiotic activated receptors or the xenosensors detect xenobiotic exposure and coordinate expression of genes involved in appropriate pharmacological or toxicological responses including antioxidant enzymes, genes involved in proliferation, drug metabolizing enzymes such as cytochrome P450 (CYP), various conjugation enzymes, and transporters [1, 2]. Nuclear receptors such as Aryl hydrocarbon Receptor (AHR), Constitutive Androstane Receptor (CAR), Pregnane X Receptor (PXR), and Peroxisome Proliferator-activated Receptors (PPARα) are the major xenosensors in the liver [3]. Upon binding to their specific ligands, these receptors bind to the DNA as heterodimers generally with RXR and activate their target genes.

Hepatocyte nuclear factor 4 alpha (HNF4α) is a highly conserved orphan nuclear receptor [4]. HNF4α is classified as an orphan receptor because it does not have a ligand that activates the receptor, which is the case for most other nuclear receptors [5]. HNF4α controls the expression of a significant number of liver-specific genes that control critical liver functions. The absence of HNF4α in the adult liver results in rapid dedifferentiation, pronounced metabolic dysregulation, hepatomegaly, spontaneous proliferation ultimately causing increased mortality [6–13]. Importantly, HNF4α regulates expression of many CYP gene involved in drug metabolism and detoxification [14]. Recent studies from our laboratory show that loss of HNF4α function is a common step in progression of chronic liver diseases including metabolism associated fatty liver disease, alcoholic liver disease, liver cirrhosis and hepatocellular carcinoma [15].

Signaling pathways and xenosensors governing xenobiotic metabolism and disposition are intertwined and crosstalk with each other [16]. Furthermore, few studies have also shown possible interaction between HNF4α and other xenosensors but the mechanisms are not completely clear. This study aimed to find the interaction between HNF4α, and AhR, CAR, PXR and PPARα. Our studies have shown that HNF4α is critical for the activation of liver-specific xenosensing receptors involved in toxicological responses.

## Materials and Methods

### Animals, Treatment, and Tissue Harvesting

All animals were housed in facilities accredited by the Association for Assessment and Accreditation of Laboratory Animal Care at the University of Kansas Medical Center under a standard 12 h light/dark cycle with free access to chow and water. All studies were approved by the Institutional Animal Care and Use Committee at the University of Kansas Medical Center. HNF4α-floxed mice were injected with AAV8-TBG-eGFP or AAV8-TBG-CRE (Vector Biolabs) intraperitoneally (i.p.) to generate wild-type (WT) and hepatocyte-specific HNF4α knockout (HNF4α-KO) mice, respectively, as described previously [10].

WT and HNF4α-KO mice were treated with mouse-specific activators of AhR, CAR, PXR, and PPARα 8 days after AAV8 injection. A single dose of 30 µg/kg of 2,3,7,8-Tetracholorodibenzo-p-dioxin (TCDD) (AccuStandard D-404S) dissolved in DMSO was injected i.p. to activate AhR. Mice were sacrificed 5 days after TCDD administration. A single dose of 2.5 µg/g of 1,4-Bis [2-(3,5-Dichloropyridyloxy)] benzene (TCPOBOP) (Millipore Sigma, T1443) dissolved in ethanol was injected to activate CAR in these mice. Mice were sacrificed 3 days after TCPOBOP treatment. WT and HNF4α-KO mice were administered a daily dose of 100 µg/g of pregnenolone 16α-carbonitrile (PCN) (Cayman Chemical, 16343) dissolved in ethanol for 4 days to activate PXR and were sacrificed on day 5. PPARα was activated in WT and HNF4α-Ko mice by injecting 1 mg/kg of WY-14643 (Selleckchem, S8029) dissolved in DMSO daily for 5 days. Mice were sacrificed on Day 6. In all experiments, control mice were treated with corn oil and vehicle (either ethanol or DMSO). After sacrificing the mice, liver and blood samples were collected and processed separately to obtain paraffin sections, frozen sections, and serum separation, as described previously [17].

### Protein isolation and western blot analysis

Frozen liver tissues were used to make liver lysates. Total protein was isolated from the livers of WT and HNF4α-KO mice using Pierce RIPA buffer (Thermo Scientific 89901) with protease and phosphatase inhibitors (Halt Protease and Phosphatase Inhibitor Cocktail with EDTA; Thermo Scientific 78438) freshly added to it at a concentration of 1:100. Cell lysates were prepared using beaded tubes. Bicinchoninic acid protein assay reagents (Thermo Fisher Scientific) were used to determine the protein concentration. Pooled samples of protein extracts (80 μg) were separated using electrophoresis on 4– 12% NuPage Bis-Tris gels with MOPS buffer (Invitrogen, Carlsbad, CA) and then transferred to Immobilon-P membranes (Millipore) in NuPAGE transfer buffer with added 20% methanol. Loading and transfer efficiency were verified by staining membranes with Ponceau S. Membranes were probed with respective primary and secondary antibodies dissolved in 5% nonfat milk or 5% bovine serum albumin in Tris-buffered saline with Tween 20. Li-COR Odyssey FC was used for imaging the western blots. The details about the antibodies used are listed in Table 1.

**Table 1:** Antibodies used in this study.

| Name | Manufacturer | Catalog number | Dilution |
| --- | --- | --- | --- |
| HNF4α | Perseus Proteomics | PP-H1415-00 | 1:2000 |
| Cyclin D1 | Cell Signaling Technology | 2978 | 1:1000 |
| GAPDH | Cell Signaling Technology | 2118S | 1:5000 |

### Real-Time Polymerase Chain Reaction

Total RNA was isolated from frozen liver tissues using the TRIzol method, according to the manufacturer’s protocol (Sigma-Aldrich). RNA was converted to cDNA using a high-capacity cDNA reverse transcription kit (Applied Biosystem 4368814) according to the manufacturer’s protocol. The mRNA levels of various genes were detected by real-time PCR analysis using the SYBR green method, as described previously in detail [18].18s was used as an internal control. Changes in the genes of interest were normalized to 18 s mRNA, and data is presented as a fold change compared with control WT mice. The primer sequences for real-time PCR are listed in Table 2.

**Table 2:**
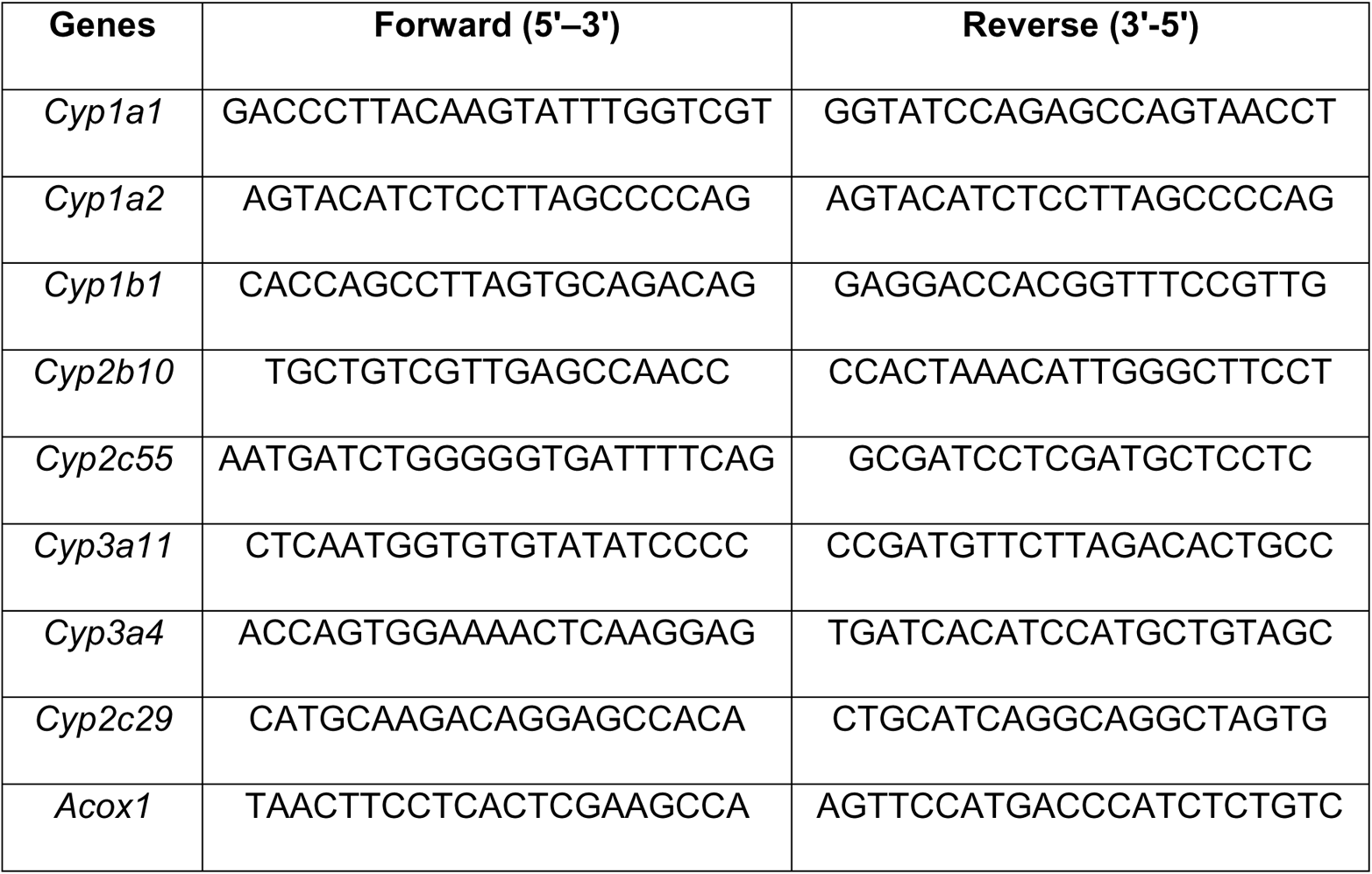
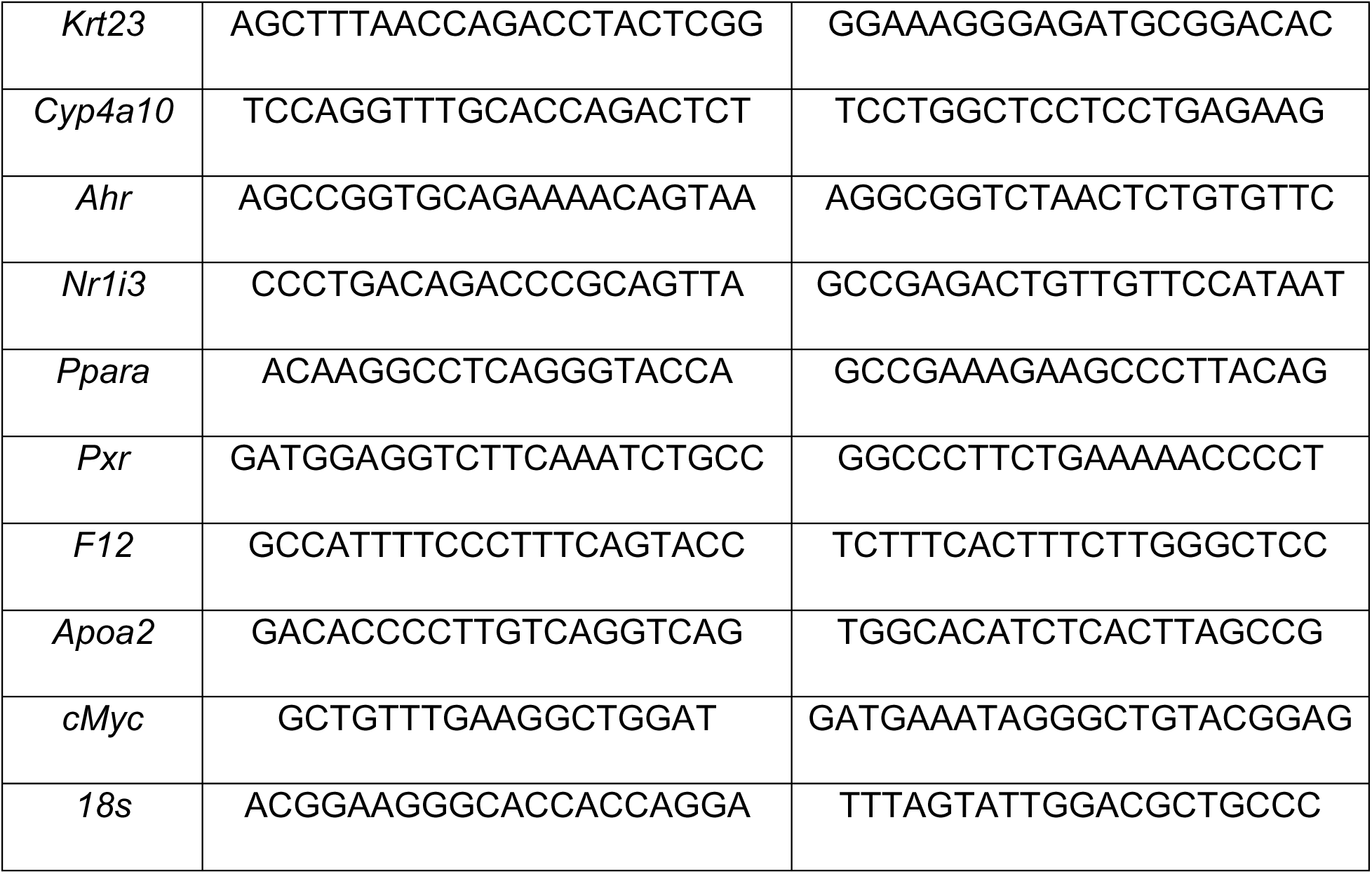
Primers used in this study.

### Staining procedures

5 μM thick paraffin-embedded liver sections were used for hematoxylin and eosin (H&E) staining and Ki67 (Cell signaling technology 12202S, 1:400) immunohistochemical staining. During immunohistochemical staining, slides were kept in boiling citrate buffer for 30 minutes to achieve antigen retrieval. 5% normal goat serum was used to block the tissue section, followed by overnight incubation with the primary antibody at 4°C. The avidin–biotin complex method (ABC Vectastain kit; Vector Laboratories, Burlingame, CA) was then used to link the primary antibody to biotinylated secondary antibody, followed by the use of chromogen diaminobenzidine to visualize the stained protein.

### Serum ALT

A Pointe Scientific ALT Assay kit by Fisher Scientific was used to measure serum alanine aminotransferase (ALT).

### Statistical analysis

Data presented in the form of bar graphs show the mean ± standard error of the mean. GraphPad Prism 9 was used to graph and calculate statistics. Two-way ANOVA and Student’s *t-test* were applied to all analyses, with *p*<0.05 considered significant.

## Results

### AhR activation after TCDD treatment

WT and HNF4α-KO mice were administered AhR agonist, TCDD, and control mice were given corn oil (Fig. 1). The HNF4α-KO mice had a significantly higher liver weight to body weight ratio (LW/BW), a finding consistent with previous studies [5, 10–12]. TCDD treatment did not affect LW/BW in either WT or HNF4α-KO mice (Fig. 2A). Serum ALT levels were increased in only HNF4α-KO mice after TCDD administration, but not significantly (Fig. 2B). Hepatic protein levels of the proliferation marker CyclinD1 were not changed in WT but decreased in HNF4α-KO mice after TCDD treatment, as confirmed by Western blot (Fig. 2C). There were no histological changes in the WT and HNF4α-KO mice after TCDD treatment (Fig. 2D). HNF4α-KO mice are known to show higher proliferation at the basal level [11]. TCDD administration did not affect the proliferation response in the WT mice but significantly reduced it in HNF4α-KO mice (Fig. 2E, 2F). AhR activation was analyzed by quantifying mRNA of its classic target genes. AhR target genes *Cyp1a1, Cyp1a2*, and *Cyp1b1* showed reduced expression in HNF4α-KO mice at the basal level. TCDD treatment significantly induced *Cyp1a1* and *Cyp1a2* in both WT and HNF4α-KO mice whereas showed no change in *Cyp1b1* expression (Fig. 2G–I).

**Figure 1:**
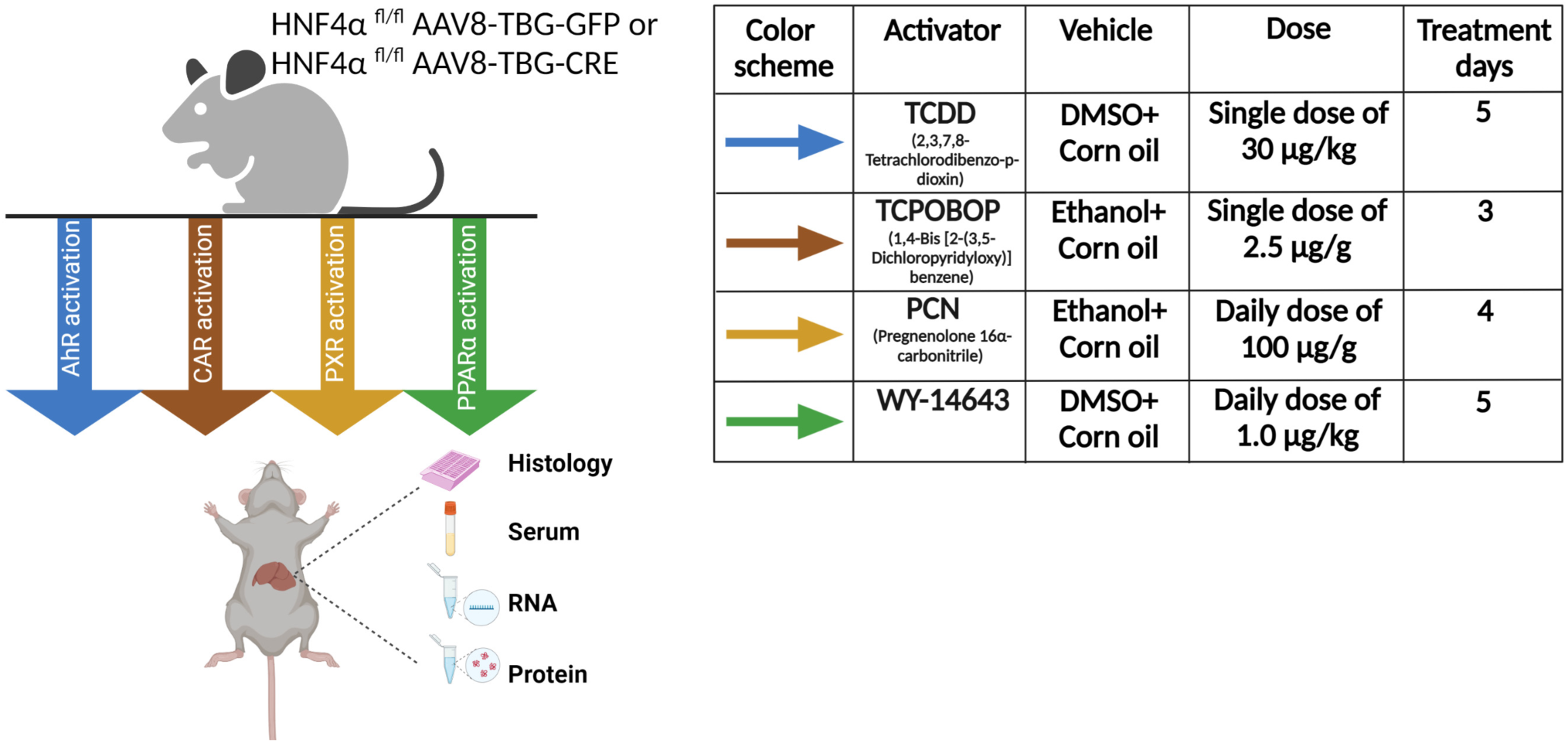
Schematic representation of the experimental design.

**Figure 2:**
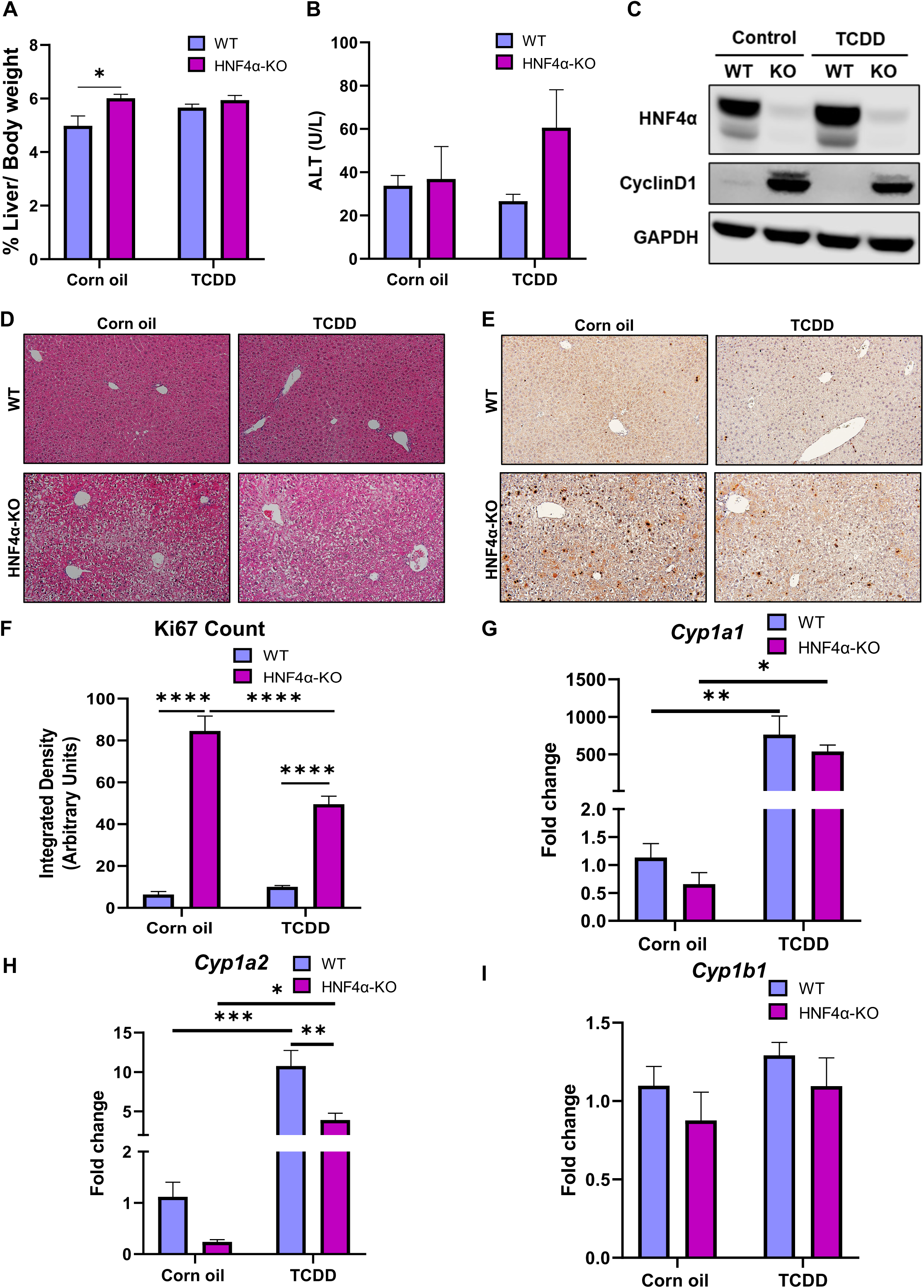
AhR activation after TCDD treatment. (A) Liver weight to body weight ratio, (B) serum ALT levels, (C) Western blot analysis of total hepatic HNF4α, cyclin D1, and GAPDH in WT and HNF4α-KO mice after corn oil and TCDD treatment. Photomicrographs of (D) H&E staining and (E) Ki67 staining and (F) quantification of Ki67 positive cells. qPCR analysis of (G) *Cyp1a1*, (H) *Cyp1a2*, and (I) *Cyp1b1* for WT and HNF4α-KO mice after corn oil and TCDD administration. Original magnification, 200X. The bar represents the mean ± standard error of the mean. *P<0.05, **P < 0.01, ***P<0.001, ****P<0.0001.

### CAR activation after TCPOBOP treatment

WT and HNF4α-KO mice were treated with TCPOBOP, a CAR agonist, and control mice were given corn oil (Fig. 1). TCPOBOP treatment significantly increased LW/BW ratio in WT mice but not in the HNF4α-KO mice (Fig. 3A). WT mice showed moderate liver injury as demonstrated by a 3-fold increase in serum ALT levels. However, serum ALT levels did not change in HNF4α-KO mice after TCPOBOP treatment (Fig. 3B). TCPOBOP administration increased hepatic protein levels of cyclin D1 in WT mice but a decrease in HNF4α-KO mice (Fig. 3C). H&E staining indicated no major histological changes in either the WT or HNF4α-KO mice after TCPOBOP administration (Fig. 3D). Significantly high Ki67-positive cells were observed in the WT mice after TCPOBOP treatment. Interestingly, TCPOBOP treatment reduced the expression of Ki67 in HNF4α-KO mice (Fig. 3E, 3F). CAR activation was analyzed by quantifying the mRNA levels of classic CAR target genes such as *Cyp3a11, Cyp2b10*, and *Cyp2c55*. These genes were significantly induced in WT mice after TCPOBOP treatment. Strikingly, TCPOBOP did not induce any of the CAR target genes in HNF4α-KO mice (Fig. 3G–I).

**Figure 3:**
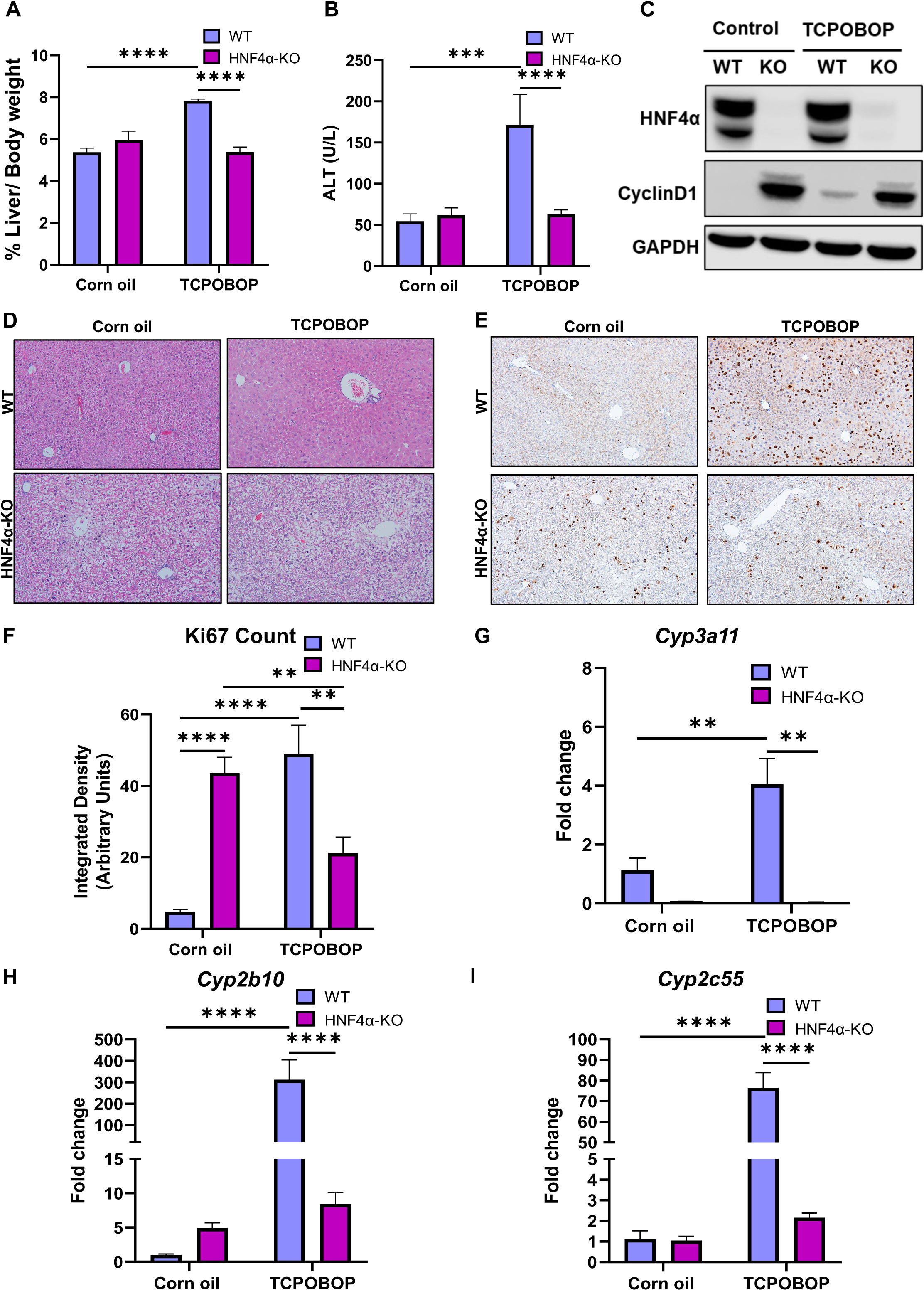
CAR activation after TCPOBOP treatment. (A) Liver weight to body weight ratio, (B) serum ALT levels, (C) Western blot analysis of total hepatic HNF4α, Cyclin D1, PCNA, and GAPDH in WT and HNF4α-KO mice after corn oil and TCPOBOP treatment. Photomicrographs of (D) H&E staining and (E) Ki67 staining and (F) quantification of Ki67 positive cells. qPCR analysis of (G) *Cyp3a11*, (H) *Cyp2b10*, and (I) *Cyp2c55* for WT and HNF4α-KO mice after corn oil and TCPOBOP administration. Original magnification, 200X. The bar represents the mean ± standard error of the mean. *P<0.05, **P < 0.01, ***P<0.001, ****P<0.0001.

### PXR activation after PCN treatment

WT and HNFα-KO mice were treated with the mouse PXR agonist, PCN, to activate PXR, and control mice were given corn oil (Fig. 1). PCN administration significantly increased LW/BW ratio in both WT and HNF4α-KO mice (Fig. 4A). PCN treatment did not affect serum ALT levels in either WT or HNF4α-KO mice (Fig. 4B). Western blot analysis showed only a slight increase in Cyclin D1 protein expression in WT mice and a further increase in already higher levels of Cyclin D1 protein in HNF4α-KO mice after PCN administration (Fig. 4C). PCN treatment did not show any major changes in liver histology in either WT or HNF4α-KO mice (Fig. 4D). HNF4α-KO mice showed higher Ki67 positive cells at basal levels, which further increased remarkably after PCN administration (Fig. 4E, 4F). Expression of PXR target genes including *Cyp3a11*, *Cyp3a4*, and *Cyp2c29* were significantly upregulated in WT mice after PCN treatment. HNF4α-KO mice showed a reduced expression of PXR target genes at the basal level and no significant increase was observed in expression of these genes in HNF4α-KO mice following PCN treatment (Fig. 4G–I).

**Figure 4:**
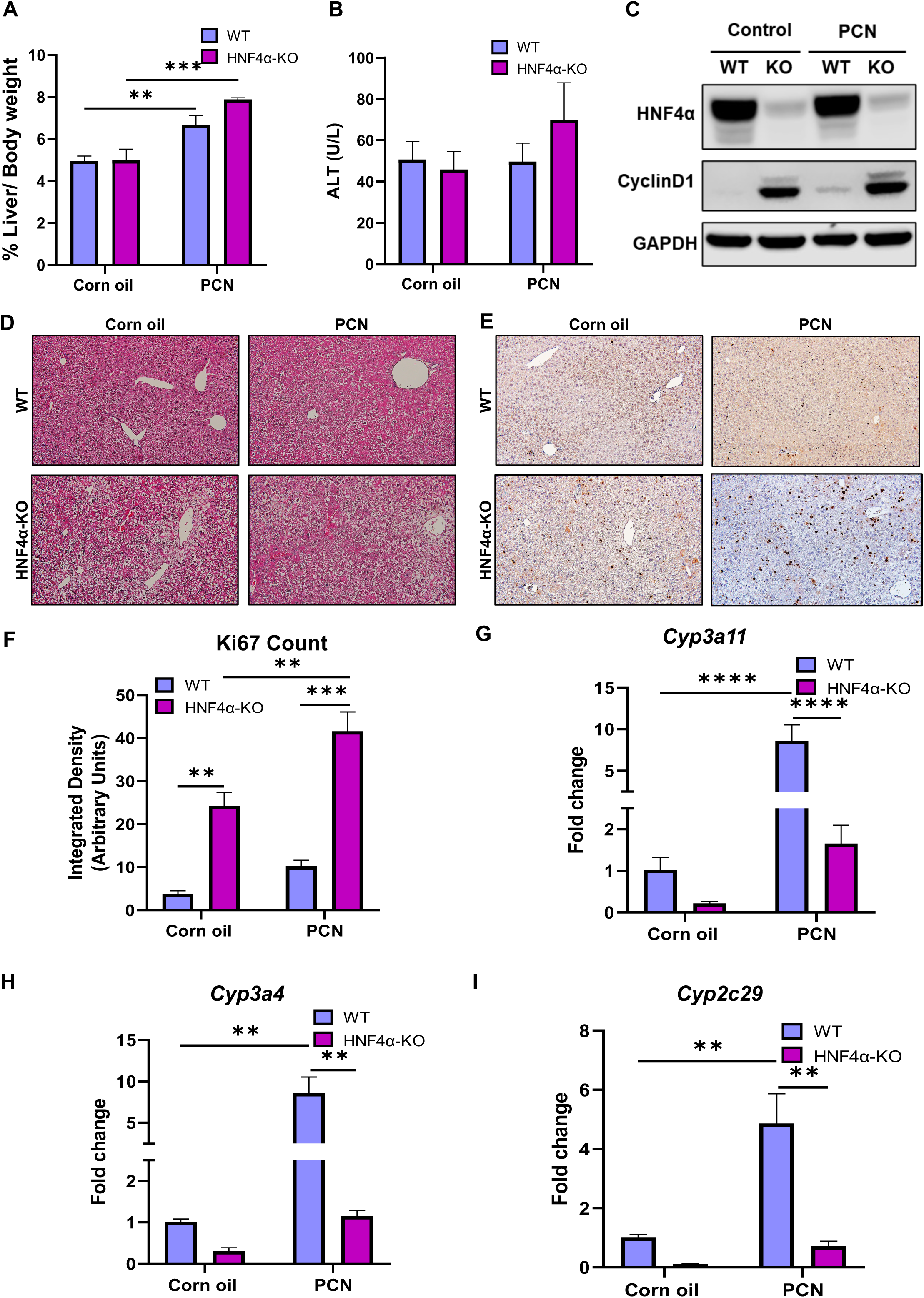
PXR activation after PCN treatment. (A) Liver weight to body weight ratio, (B) serum ALT levels, (C) Western blot analysis of total hepatic HNF4α, Cyclin D1, PCNA, and GAPDH in WT and HNF4α-KO mice after corn oil and PCN treatment. Photomicrographs of (D) H&E staining and (E) Ki67 staining and (F) quantification of Ki67 positive cells. qPCR analysis of (G) *Cyp3a11*, (H) *Cyp3a4*, and (I) *Cyp2c29* for WT and HNF4α-KO mice after corn oil and PCN administration. Original magnification, 200X. The bar represents the mean ± standard error of the mean. *P<0.05, **P < 0.01, ***P<0.001, ****P<0.0001.

### PPARα activation after WY-14643 treatment

WT and HNF4α-KO mice were administered PPARα agonist, WY-14643 and control mice were administered corn oil (Fig. 1). WY-14643 treatment resulted in a significantly increased LW/BW ratio in HNF4α-KO mice, but it remained unchanged in WT mice (Fig. 5A). There was no change in serum ALT levels in either WT or HNF4α-KO mice after WY-14643 administration (Fig. 5B). Protein levels of cyclin D1 were reduced in HNF4α-KO mice after WY-14643 treatment (Fig. 5C). WY-14643 treatment did not cause any histological changes in either WT or HNF4α-KO mice (Fig. 5D). WY-14643 administration did not change Ki67 protein expression in the WT mice and it failed to induce any further increase in the ongoing proliferation in the HNF4α-KO mice (Fig. 5E, 5F). Activation of PPARα target genes including *Acox1, Krt23*, and *Cyp4a10* was analyzed following WY14643 treatment, which showed significant upregulation in WT mice. Interestingly, all PPARα target genes showed reduced expression in HNF4α-KO mice at the basal level and there was no change in the expression of these genes after WY-14643 treatment (Fig. 5G–I).

**Figure 5:**
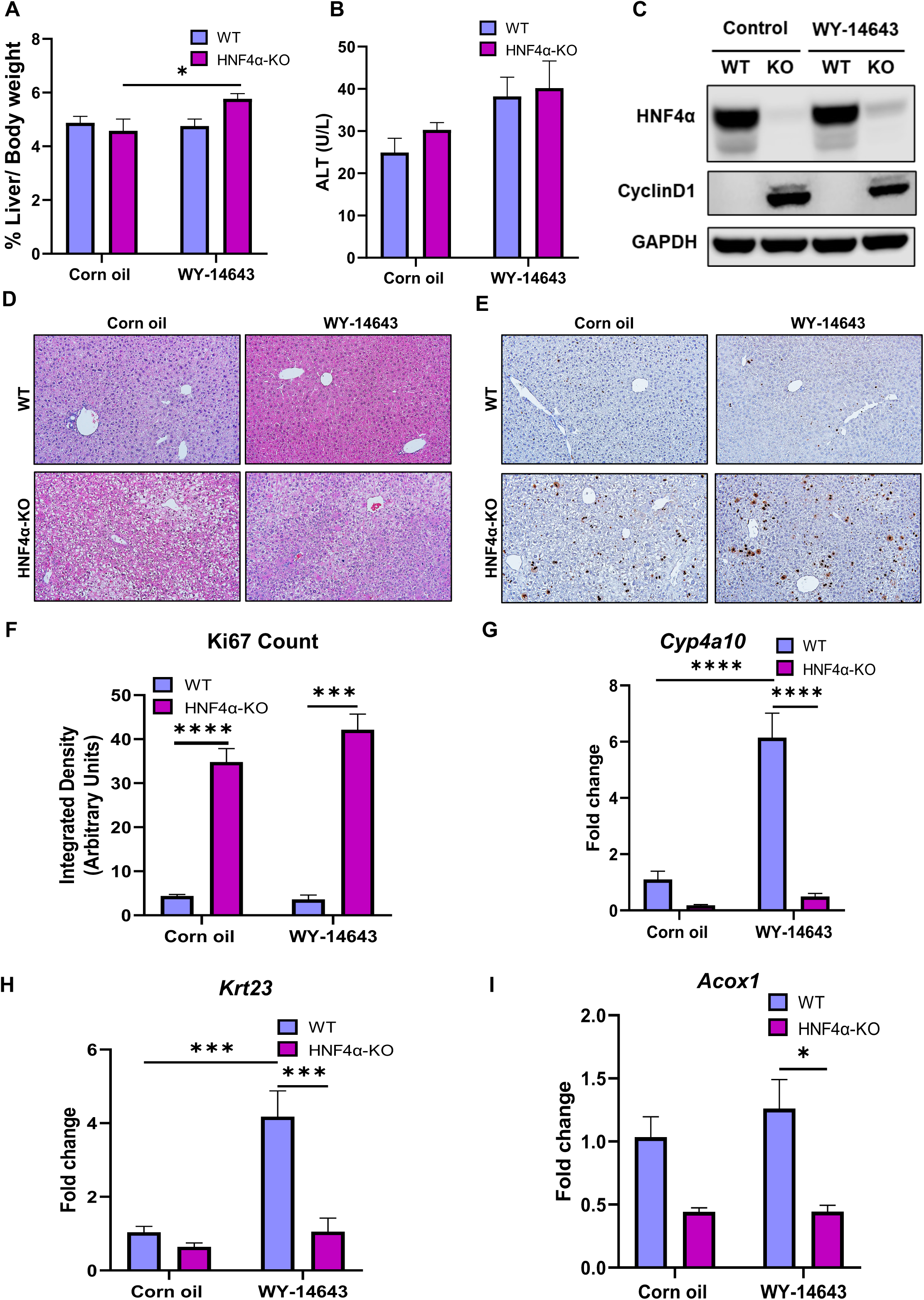
PPARα activation after WY-14643 treatment. (A) Liver weight to body weight ratio (B) Serum ALT levels of WT and HNF4α-KO mice after corn oil and WY-14643 treatment. (C) Western blot analysis of total hepatic HNF4α, Cyclin D1, PCNA, and GAPDH in WT and HNF4α-KO mice after corn oil and WY-14643 treatment. Photomicrographs of (D) H&E staining and (E) Ki67 staining and (F) quantification of Ki67 positive cells. qPCR analysis of (G) *Cyp4a10*, (H) *Krt23*, and (I) *Acox1* for WT and HNF4α-KO mice after corn oil and PCN administration. Original magnification, 200X. The bar represents the mean ± standard error of the mean. *P<0.05, **P < 0.01, ***P<0.001, ****P<0.0001.

### Cross talk between HNF4α and xenosensors

To determine the effect of HNF4α deletion on the gene expression of xenosensors, qPCR analysis for *Ahr, Nr1i3* (gene for CAR), *Ppara*, and *Nr1i2* (gene for PXR) was performed with control WT and HNF4α-KO mice. We observed that *Ahr, Nr1i3*, and *Ppara* transcript levels were significantly reduced upon deletion of HNF4α, whereas *Nr1i2* was upregulated in HNF4α-KO mice as compared to WT mice (Fig. 6).

**Figure 6:**
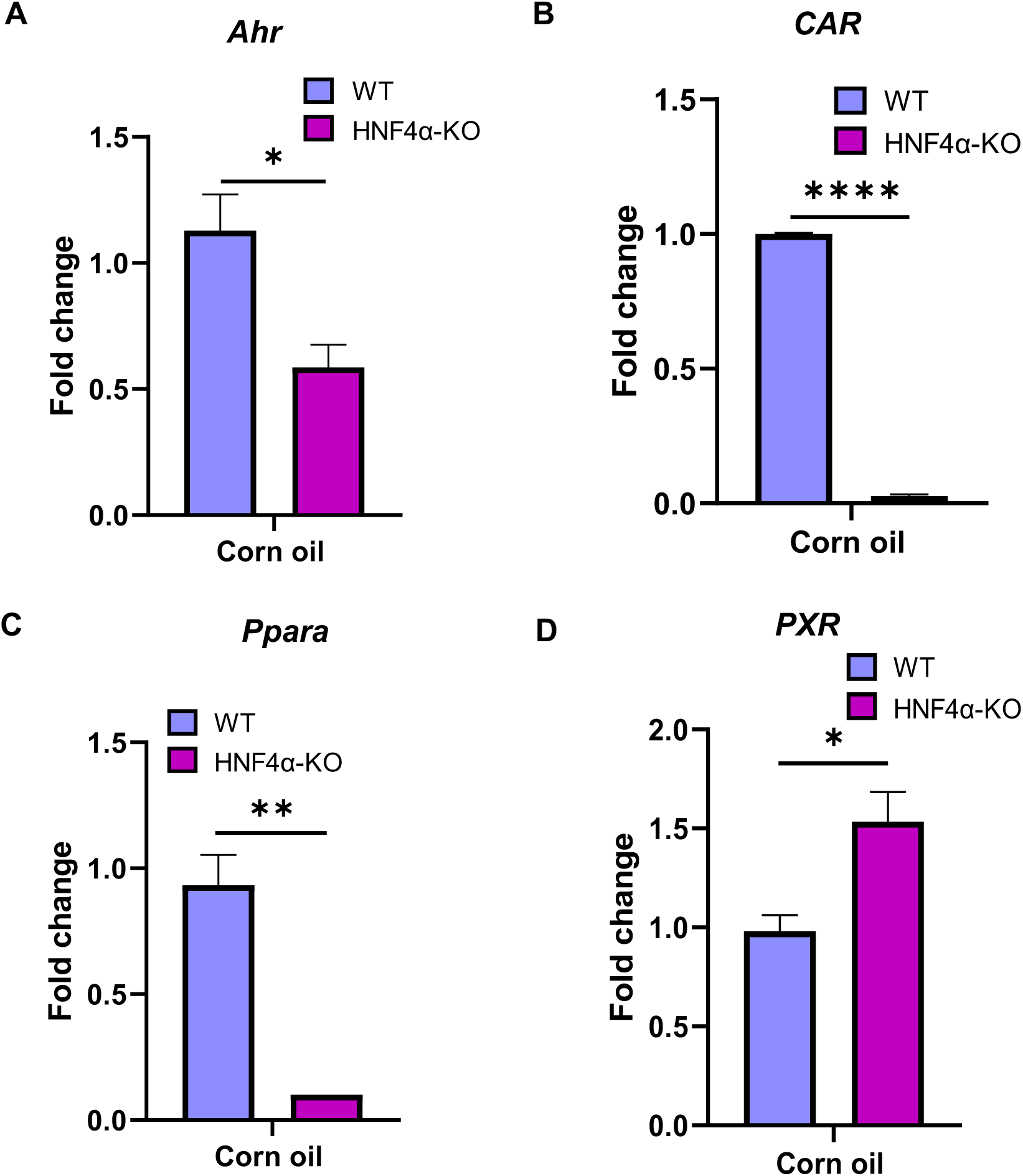
qPCR analysis of (A) *Ahr*, (B) *Nr1i3r*, (C) *Nr1i2* and (D) *Ppara* in control WT and HNF4α-KO mice treated with corn oil. The bar represents the mean ± standard error of the mean. *P<0.05, **P < 0.01, ****P<0.0001.

Further, we quantified the mRNA expression of HNF4α target genes, *F12* and *Apoa2.* These genes were significantly induced in WT mice at the basal level. TCDD treatment, AhR activation, did not change the expression of these genes in either WT or HNF4α-KO mice. TCPOBOP (CAR activation), PCN (PXR activation), and WY-14643 (PPARα activation) treatments significantly decreased the expression of *F12* and *Apoa2* in WT mice as compared to the control WT mice (Fig. 7).

**Figure 7:**
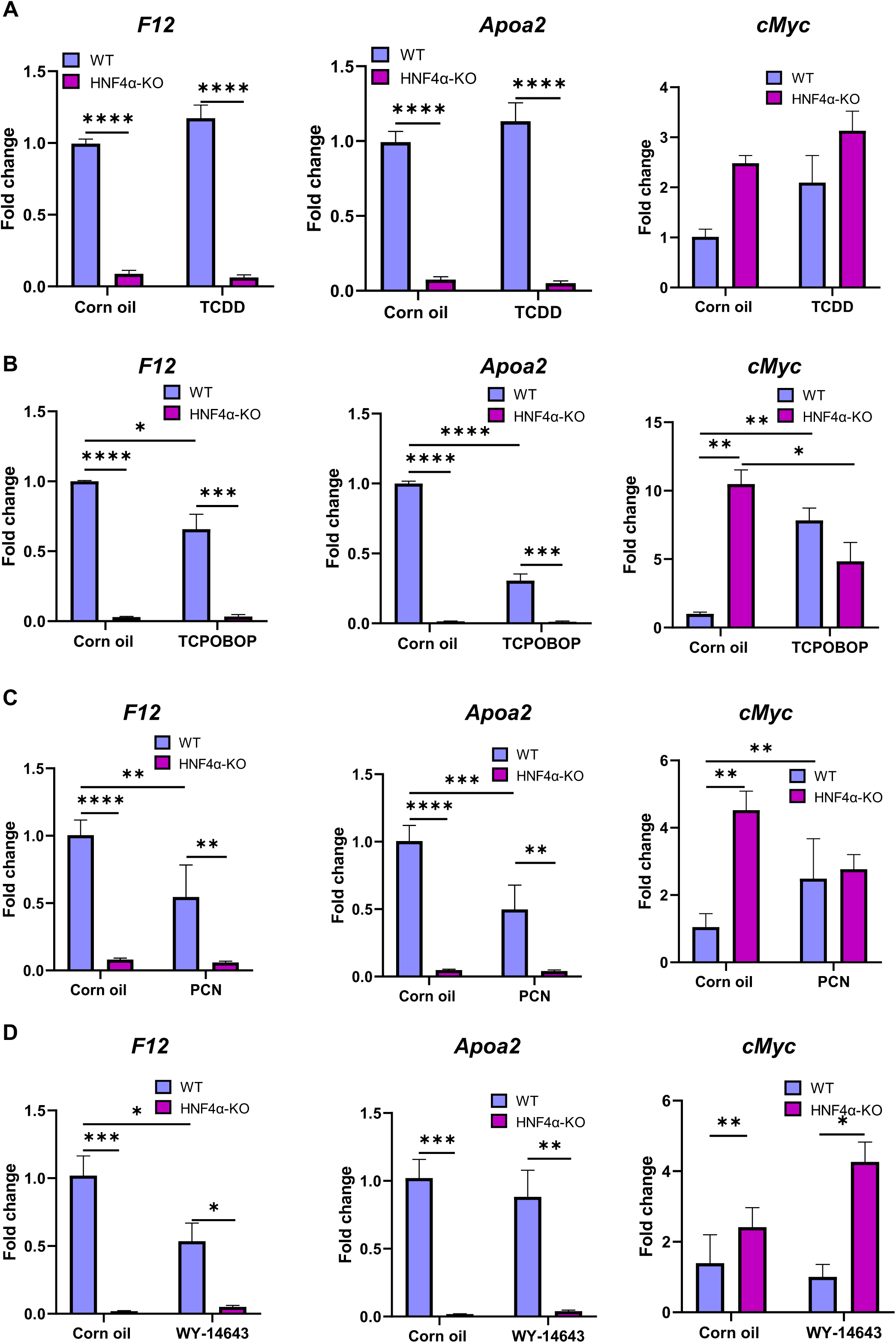
qPCR analysis of genes positively regulated by HNF4α including *F12* and *Apoa2* in WT and HNF4α-KO mice after (A) TCDD treatment, (B) TCPOBOP treatment, (C) PCN treatment, and (D) WY-14643 treatment.

## Discussion

HNF4α is a key nuclear receptor in the liver expressed in hepatocytes. It plays an important role during embryonic and postnatal liver development. HNF4α is critical for the maintenance of differentiated phenotypes, as it regulates the expression of a significant number of hepatocyte-specific genes involved in nutrient metabolism, gluconeogenesis, lipid metabolism, bile acid biosynthesis, blood coagulation and drug metabolism, and detoxification enzymes (DMEs) [11, 14, 19, 20]. The liver detoxifies and facilitates the excretion of xenobiotics by converting lipid-soluble compounds to more water-soluble compounds enzymatically with the help of DMEs, including CYP [21]. HNF4α is a regulator that controls the expression of several major drug-metabolizing CYP [22].

The expression of CYP enzymes is upregulated in response to exposure to high concentrations of xenobiotics. The induction of CYP is also regulated by ligand-activated nuclear receptors, including AhR, CAR, PXR and PPARα. These nuclear receptors function as xenosensors because they detect xenobiotic exposure in the system and regulate responses to such exposures [3]. Along with many endogenous and exogenous compounds, environmental chemicals, such as polycyclic aromatic hydrocarbons, polychlorinated biphenyls, and fibrates, are known ligands for xenosensors. AhR is a transcription factor and member of a Per/ARNT/Sim (PAS) family that coordinates the regulation of CYP1 enzymes. AhR ligands are environmental pollutants of polyhalogenated aromatic hydrocarbons. Along with drug metabolism, AhR also regulates the many genes involved in immune response and energy homeostasis [23]. CAR and PXR are the key xenosensors that activate the hepatic expression of the majority of DMEs, including CYP2B, 2C, and 3A [24]. The hepatic expression of CAR and PXR is transactivated by HNF4α [25, 26]. In addition to the regulation of DMEs, CAR regulates the expression of genes involved in gluconeogenesis and lipogenesis [27]. PPARα is an important regulator of the hepatic metabolism of drugs and lipids. The prompter of PPARα is activated by HNF4α in addition to PPARα itself [28]. PPARα is known to induce CYP4 enzymes.

Many of the drug detoxifications and metabolic processes are regulated by both HNF4α and these major nuclear receptors. AhR, CAR, PXR and PPARα interact with HNF4α to regulate the expression of DMEs [29]. However, the crosstalk between HNF4α and xenosensors is not well established. Previous studies done in context with HNF4α are mostly in vitro using hepatic cell lines. Very little is known about the activation of xenosensors in the absence of HNF4α. The goal of this study was to investigate the role of HNF4α in the activation of hepatic xenosensors. We hypothesize that HNF4α is essential for the function of these four major xenobiotic-sensing nuclear receptors.

In this study, WT and HNF4α-KO mice were administered mouse-specific activators of xenosensors for acute activation. We focused on four major aspects such as LW/BW ratio, liver injury, proliferation, and mRNA expression of target genes of the nuclear receptors of interest—to study their activation. In our first experiment, AhR activation, there was no significant increase in liver injury in either WT or HNF4α-KO mice. Loss of HNF4α is known to increase the proliferation response [30]. AhR is known to positively regulate cell proliferation and survival via regulating cell cycle entry, growth factor signaling, receptor expression, and apoptosis [31]. In our study, there was no effect on the proliferation response in WT mice after acute activation of AhR. However, AhR activation significantly lowered proliferation in HNF4α-KO mice. Further, deletion of HNF4α resulted in a significant reduction of *Ahr* mRNA levels at the basal level. The induction of the classic AhR target genes *Cyp1a1* and *Cyp1a2* in both WT and HNF4α-KO mice after TCDD treatment indicated that AhR activation was not affected in the absence of HNF4α.

CAR is a key member of the xenosensor family, which regulates the expression of the most important CYP enzymes involved in the detoxification response. TCPOBOP is a mouse-specific agonist of CAR. Acute exposure to xenobiotics has been shown to induce CAR-mediated hepatomegaly and an increased proliferation response in mice [32]. Consistent with these results, in the experiment of CAR activation, we found that TCPOBOP administration significantly increased LW/BW ratio, liver injury, and proliferation in WT mice. Interestingly, TCPOBOP induced hepatomegaly, and injury was absent in HNF4α-KO mice. In addition, TCPOBOP treatment resulted in a significantly reduced proliferation response in HNF4α-KO mice. In WT mice, CAR target genes were significantly induced after TCPOBOP treatment, which confirmed CAR activation in these mice. However, in HNF4α-KO mice, the expression of these genes was low at the basal level, and TCPOBOP treatment did not induce them. Previous studies have shown that basal CAR expression is diminished in HNF4α-KO mice [25]. Our studies were consistent with these observations. The mRNA expression of *Nr1i3* (gene for CAR) was significantly low in the absence of HNF4α. These results indicate that CAR activation was disrupted in the absence of HNF4α.

PXR is a key regulator of xenobiotic metabolism. It regulates the expression of several important CYPs, including CYP3. PCN is a mouse-specific ligand that activates PXR. PXR activation is known to induce hepatomegaly [33]. Consistent with these findings, we observed a significant increase in LW/BW ratio after PCN administration in WT as well as HNF4α-Ko mice. PXR activation also affects the proliferation response. Studies have shown that PXR activation is important in driving cell proliferation during the regeneration phase after partial hepatectomy [34]. In our study, PXR activation showed a slight increase in the proliferation response in WT mice. However, the proliferation response induced by the deletion of HNF4α was further doubled after PCN administration in HNF4α-KO mice. PXR activation, analyzed by mRNA expression of PXR target genes, was confirmed in the WT mice. Interestingly, the low basal levels of these genes in HNF4α-KO were not changed after PCN treatment, indicating that PXR was not activated in the absence of HNF4α. However, we observed increased mRNA expression of *Nr1i2* (a gene for PXR) in HNF4α-KO mice at the basal level.

Two weeks of WY-14643 treatment has been shown to induce hepatomegaly and increase cell proliferation in WT [35]. WY-14643 is known to induce cell proliferation in mouse livers after long-term exposure [36]. However, in our studies, we did not see any change in LW/BW ratio and proliferation response in WT mice administered WY-14643. We think it is because of the acute exposure to WY-14643 that we did not see any difference in cell proliferation. In contrast, we observed an increased LW/BW ratio in HNF4α-KO mice after the treatment. HNF4α is shown to transcriptionally regulate PPARα [37]. Consistent with these results, we observed that *Ppara* gene expression was significantly reduced in HNF4α-KO mice at the basal level. PPARα target gene expression was not upregulated in the absence of HNF4α specifying that PPARα activation was disrupted in these mice.

In our next experiment, we observed that genes positively regulated by HNF4α including *F12* and *Apoa2*, were significantly downregulated in WT mice after CAR, PXR, and PPARα activation. There was no change in the expression of these genes after AhR activation. This suggests that not AhR activation but CAR, PXR, and PPARα activation affects HNF4α activation. A recent publication from our laboratory has shown that the progressive loss of HNF4α is associated with chronic liver disease progression [15]. The results from the current study showed that in the absence of HNF4α, toxicologically important nuclear receptor activation is disrupted. Xenosensors orchestrate the regulation of CYPs, which act on common enzymatic pathways by numerous drugs; therefore, maintaining xenosensor activation and function is pivotal in disease conditions [38]. Disrupted activation of xenosensors during disease conditions might result in poor induction of genes in the disposition of pharmacological agents and environmental toxicants. Therefore, maintaining HNF4α function in disease conditions is necessary to activate xenobiotic sensors.

In summary, our studies show that the xenosensor’s mechanism of regulating proliferation is different from xenobiotic metabolism. Deletion of HNF4α significantly reduced the expression of CAR, PXR, and PPARα target genes; however, the AhR target genes were not affected. In conclusion, HNF4α is required for CAR, PXR, and PPARα function and is important for pharmacological and toxicological responses governed by major hepatic xenosensors (Fig. 8).

**Figure 8:**
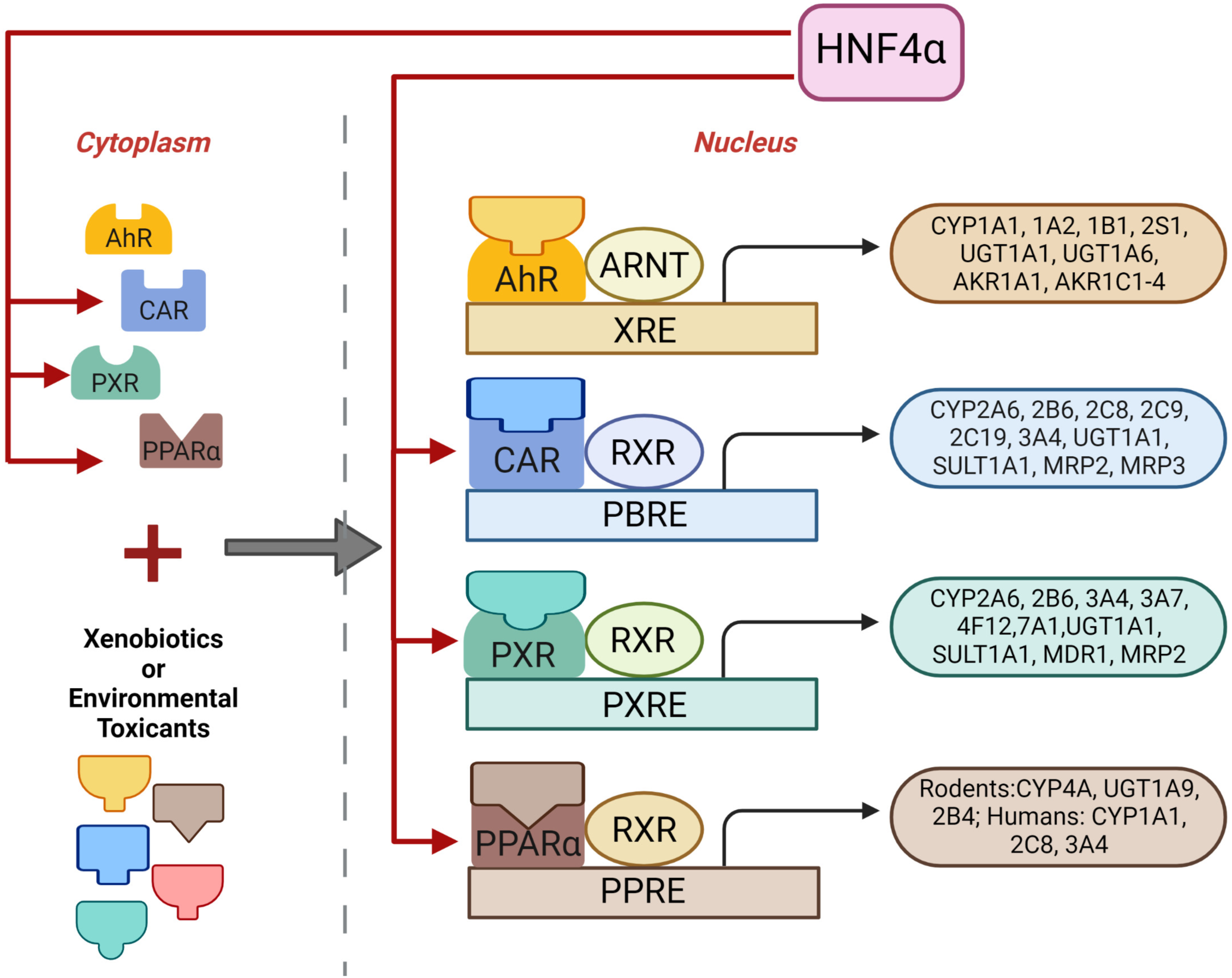
Regulation hepatic xenosensor function by HNF4α.

